# kmindex and ORA: indexing and real-time user-friendly queries in terabyte-sized complex genomic datasets

**DOI:** 10.1101/2023.05.31.543043

**Authors:** Téo Lemane, Nolan Lezzoche, Julien Lecubin, Eric Pelletier, Magali Lescot, Rayan Chikhi, Pierre Peterlongo

## Abstract

Public sequencing databases contain vast amounts of biological information, yet they are largely underutilized as one cannot efficiently search them for any sequence(s) of interest. We present `kmindex`, an innovative approach that can index thousands of highly complex metagenomes and perform sequence searches in a fraction of a second. The index construction is an order of magnitude faster than previous methods, while search times are two orders of magnitude faster. With negligible false positive rates below 0.01%, `kmindex` outperforms the precision of existing approaches by four orders of magnitude. We demonstrate the scalability of `kmindex` by successfully indexing 1,393 complex marine seawater metagenome samples from the *Tara* Oceans project. Additionally, we introduce the publicly accessible web server “Ocean Read Atlas” (`ORA`) at https://ocean-read-atlas.mio.osupytheas.fr/, which enables real-time queries on the *Tara* Oceans dataset. The open-source `kmindex` software is available at https://github.com/tlemane/kmindex.

---

Public genomic datasets are growing at an exponential rate [8]. They contain treasures of genomic information that enable groundbreaking discoveries in fundamental domains such as agronomy, ecology, and health [10, 19]. Unfortunately, despite their public availability in repositories such as the Sequence Read Archive [14], these resources, measured in petabytes, are rarely ever reused globally because they cannot be searched. Recent years have seen many methodological developments towards sequencing data search engines (see [5] and [16] for a review).

Current methods for searching genomic sequencing data look for *k*-mers (words of fixed length *k*, with *k* usually in [20; 50]) shared between a query sequence and each sample present in a reference database. The central operation is thus to determine, for each *k*-mer, in which indexed sample(s) it occurs.

In this work, we focus on the challenge of indexing and querying large and complex metagenomic datasets. Given the data size, i.e. thousands of samples totalling tens of terabytes of compressed data, and its complexity, i.e. thousands of billions of distinct *k*-mers, the computational challenge is immense. Once *k*-mers are extracted from raw data and filtered, a data structure is built to associate each *k*-mer to the sample(s) in which it occurs.

Techniques for association *k*-mers to samples can be divided into three categories: 1. sketching approaches that heavily subsample *k*-mers; 2. exact data structures storing all *k*-mers; 3. approximate membership data structures e.g. Bloom filters. Sketching approaches such as sourmash [20], or Needle [9] typically suffer from high false negative rates when short sequences are queried, and are thus out of the scope of this work. Methods based on exact representations (e.g. `MetaGraph` [13], `BiFrost` [12], and `ggcat` [6]) suffer from low scalability, as highlighted by our results. We are thus left with methods based on Bloom filters (BFs) [4], such as `COBS` [3], `SBT` [21], later improved by `HowDeSBT` [11], and more recently by `MetaProFi` [22] able to index billions of *k*-mers using only a few dozen of gigabytes of space.

When indexing large and complex metagenomic datasets, existing tools face significant limitations in either 1. disk usage; 2. memory usage; 3. computation time (either during indexing and/or query); 4. false positive rate; and 5. false negative rate. Overcoming simultaneously all these limitations makes the design of an efficient data indexing strategy particularly challenging. We present `kmindex`, a new tool that performs indexing and queries using orders of magnitudes less resources than previous approaches. Also, `kmindex` provides results with no false negative calls and with negligible false positive (FP) rates, approximately four orders of magnitude smaller than those obtained by other tools. `kmindex` is primarily designed for indexing complex sequencing samples. Due to engineering choices, it is currently not suited for indexing large collections of genomes (i.e. hundreds of thousands of samples).

To showcase the features of `kmindex` on a dataset of high biological interest, we introduce a web server named “Ocean Read Atlas” (ORA) available at this URL: https://ocean-read-atlas.mio.osupytheas.fr/. `ORA` allows to search one or several sequences across all of *Tara* Oceans metagenomic raw sequencing data [23]. It enables the visualization of the results on a geographic map and their correlation with each of the 56 environmental variables collected during the circumnavigation campaign. The ORA server enables to perform instant searches on a large and complex dataset, providing new perspectives on the deep exploitation of *Tara* Oceans resources.

## Results

### Comparative results indexing 50 metagenomic seawater samples

We evaluated the performances of `kmindex` together with eight state-of-the-art *k*-mer indexers: `themisto` [2]; `ggcat` [7]; `HIBF` [18]; `PAC` [17]; `MetaProFi` [22]; `MetaGraph` [13]; `Bifrost` [12]; and `COBS` [3]. The dataset for this benchmark is composed of metagenomic seawater sequencing data from 50 *Tara* Oceans samples, of 1.4TB of gzipped fastq files. It contains approximately 1,420 billion *k*-mers. Among them, approximately 394 billion are distinct, and 132 billion occur twice or more. The exhaustive list of tool versions and commands used are proposed in a companion website https://github.com/pierrepeterlongo/kmindex_benchmarks, that also reports the False Positive computation protocols and a detailed description of the dataset considered for this benchmark.

The benchmarking setup is described in Supplementary Materials. The extensive results of all the following claims are proposed in the Supplementary Materials.

#### kmindex has superior index construction performance

Among the nine tested tools, only MetaProFi, `COBS`, and `kmindex` completed the index creation phase and were able to perform queries correctly. As shown in Table 1, building an index with `kmindex` is an order of magnitude faster than `MetaProFi` and `COBS`, and uses 2.6x less memory and 6.5x less disk. The final index sizes are all within the same magnitude range, with the smallest one produced by kmindex. The `kmindex` construction took less than 3 hours, a peak RAM of 107GB, and a peak disk usage of 878GB.

**Table 1:** Overview of index construction and read query performance of `kmindex` compared to `MetaProFi` and `COBS`, on 50 *Tara* Ocean samples. These are the only tools that succeeded in building an index and perform queries. “RAM” and “Disk” columns provide the peak usage during the building process. `COBS` and `MetaProFi` RAM and disk peaks are identical as they correspond to the same *k*-mer counting and filtration step. Queries are composed of one read and 10 million reads uniformly sampled from the 50 *Tara* Oceans datasets. All executions were performed on a cold cache. Extended results are proposed in the Supplementary Materials.

|  | Build index |  |  |  | Query time |  | FP rate (%) |  |
| --- | --- | --- | --- | --- | --- | --- | --- | --- |
|  | Time | RAM<br>GB | Disk<br>GB | Index size<br>GB | Nb. queried reads<br>1 | 10 million | Average | Max |
| **MetaProFi** | 30h15 | 278 | 5,684 | 226 | 12s72 | 1h29 | 11.18 | 21.55 |
| **COBS** | 26h30 | 278 | 5,684 | 184 | 1s51 | 15h56 | 13.29 | 24.60 |
| **kmindex** | **2h56** | **107** | **878** | **164** | **0s06** | **4m21s** | **0.006** | **0.18** |

#### kmindex enables real-time queries

As shown in Table 1, at query time, `kmindex` outperforms `MetaProFi` and `COBS`, both in terms of computation time (and memory resources, see Supp Matt). `kmindex` is between 20 and 200 times faster than `MetaProFi` and `COBS` for querying one read or millions of reads. `kmindex` is capable of performing millions of queries in a matter of minutes while allowing real-time resolution for small queries. This opens the doors to analyzing complete read sets as queries, and the deployment of real-time query servers as presented in the next section. Of note, `kmindex` also offers a “fast-mode”, presented in the Supplementary Materials, that uses more RAM to achieve even faster queries.

#### kmindex allows highly accurate queries

The `kmindex`, `MetaProFi`, and `COBS` scalability is achieved thanks to the usage of Bloom Filters that generate False Positive (FP) calls at query time. FP rate analyses, summarized Table 1, show that for a similar index size, `MetaProFi` and `COBS` present sensible false positive hits, on average 11.18% and 13.29% respectively over the 50 answers (one per indexed sample). In contrast, the `kmindex` FP rate is negligible (below 10^*−*2^% on average).

### Indexing 1,393 *Tara* Oceans samples in the Ocean Read Atlas web server

Thanks to these novel possibilities offered by `kmindex`, we built and made available a public web interface able to perform queries on a dataset composed of 1,393 samples (distinct locations and distinct fraction sizes) of the *Tara* Oceans project [23] representing 36.7 TB of raw fastq.gz files. A user can query sequences, determining their similarity with the 1,393 indexed samples. A world map depicts the resulting biogeography, as well as the environmental parameters associated with the sequences.

Note that for reasons of robustness and continuity of service, the index is deployed on a networked and redundant filesystem with lower performances compared to the benchmark environment, although suitable for this type of service. Details about indexed read sets, and more information about the server architecture and setup are provided in Supplementary Materials.

The resulting web server named “Ocean Read Atlas” (`ORA`), whose representation is provided in Figure 1, extends the “Ocean Gene Atlas” server (OGA) [25, 24] that supports queries to assembled genes from *Tara* Oceans [23] and Malaspina [1]. We believe this server will be of great importance to the *Tara* Oceans consortium as a whole, and more broadly to anybody interested in marine genetic data.

**Figure 1:**
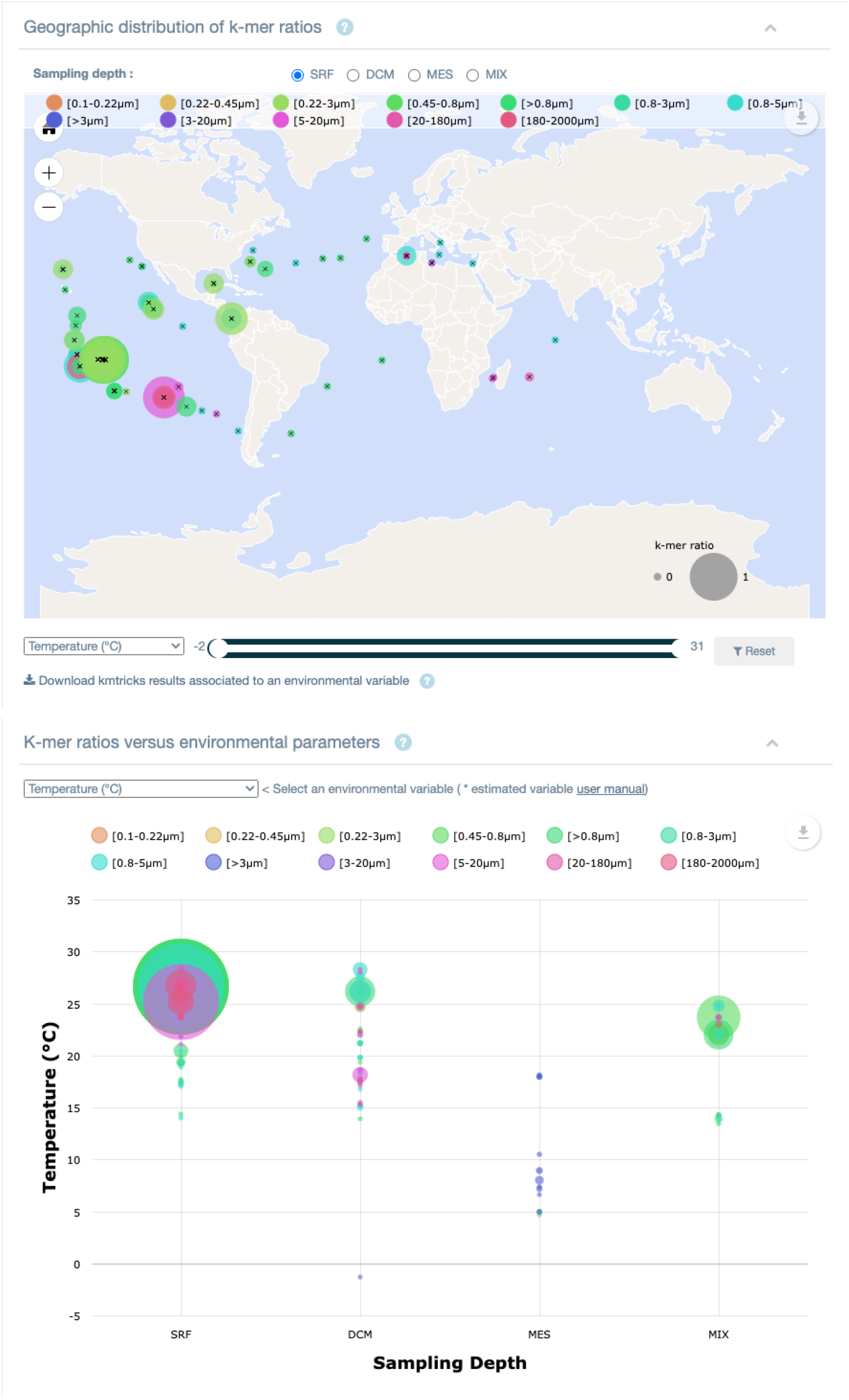
Screenshot of the “Ocean Read Atlas” result interface. Top: the biogeography distribution of the queried sequence is shown among all data samples. The size of the point depicts the similarity of the queried sequences with the corresponding sample. Bottom: a bubble plot representing the correlation between the query presence and the environmental variables of the samples in which it occurs.

#### kmindex **provides a high level of usability**

##### Index dynamicity

`kmindex` enables to add new samples to an index. A novel and independent index can be *registered* with a previous one. At query time, each registered index is queried independently. This offers the possibility to query only a subset of the registered indexes. This is well adapted when indexing samples with distinct characteristics. Alternatively, users can extend an existing index, and the parameters of the previous index (such as the *ad-hoc* hash function or the BFs sizes) are automatically reused. This second choice is less flexible but provides better performances at query time (see results presented in Supplementary Materials).

##### Versatile *k*-mer filtration

`kmindex` enables the filtration of erroneous *k*-mers, not only relying on their abundance in a dataset but also on their co-abundances in all indexed datasets. This enables to “rescue” low-abundance *k*-mers that would have otherwise been removed. To the best of our knowledge, no other indexing tool can integrate this feature. This feature is inherited from the `kmtricks` [15] algorithm.

##### Variable query resolution

`kmindex` query results can be provided with various degrees of precision. For each indexed sample, users can access the average similarity of queried sequences or a similarity value per queried sequence. Finally, `kmindex` can provide the distribution of hits, enabling to highlight some regions of interest among the queried sequences.

##### High accessibility

`kmindex` is well documented and simple to install. Queries can be performed via a CLI, via an API, or as an HTTP server.

## Methods

We briefly sum up here the `kmindex` method, while the complete description is provided in the Supplementary Materials. Conceptually, the presence of each indexed *k*-mer is stored in one BF per input read set. The BFs construction relies on kmtricks [15] which allows to filter erroneous *k*-mers and to efficiently build a partitioned matrix of BFs. Each partition indexes a subset of *k*-mers corresponding to a specific set of minimizers. In practice, with `kmindex`, matrices are *inverted* to limit cache misses during the query process, i.e. each row is a bit vector representing the presence/absence of a *k*-mer in each indexed sample. At query time, *k*-mers from queried sequences are grouped into batches, and, avoiding cache misses, BFs are queried to determine the presence or absence of each *k*-mer in each input dataset.

## Conclusion

We propose `kmindex`, a tool for creating *k*-mer indexes from terabyte-sized raw sequencing datasets. It is the only tool able to index highly complex data such as thousands of seawater metagenomic samples, and to provide instant query answers, with a non-zero but negligible false positive rate, in average below 0.01% in our tests. By its performance and its usage simplicity, `kmindex` makes indexing *k*-mers from large and complex genomic projects practically possible for the first time.

We believe that `kmindex` opens up a new channel for leveraging genetic data, removing the obstacles that often isolate studies from each other. The “Ocean Read Atlas” illustrates this advance, by providing a highly usable tool to fully utilize the wealth of data generated by the *Tara* Oceans project.

## Data Availability

A list of publicly available data used in this work is proposed in the https://github.com/pierrepeterlongo/kmindex_benchmarks repository.

## Acknowledgements

We acknowledge the GenOuest core facility (https://www.genouest.org) and the TGCC (https://www-hpc.cea.fr/index-en.html) for providing the computing infrastructure, as well as France Génomique for funding of the TGCC computing resources used to process data used in this article. The authors thank Jean-Marc Aury for his help regarding the usage of the *Tara* Oceans data sets. *Tara* Oceans (which includes both the *Tara* Oceans and *Tara* Oceans Polar Circle expeditions) would not exist without the leadership of the *Tara* Ocean Foundation and the continuous support of *Tara* Oceans consortium members. The authors also thank Kahles Andre and Mustafa Harun for their help regarding the usage of `MetaGraph`, Andrea Cracco and Alexandru Tomescu for their help using `ggcat`, and Camille Marchet and Antoine Limasset for their support using PAC. The web server is hosted by the OSU Pythéas cluster with the help of Cyrille Blanpain and SIP members. Adrien Malgoyre from SIP is thanked for the development of the OSU Pythéas GitLab.

