## Supplementary materials for "kmindex and ORA: indexing and real-time user-friendly queries in terabyte-sized complex genomic datasets"

### `kmindex` and `ORA`: Supplementary Materials

#### 1 Methods

##### 1.1 Index construction

The construction of Bloom filters from raw sequencing data is delegated to `kmtricks` [7], which allows partitioned construction of one-hash Bloom filter matrices. For each input dataset, $k$-mers are counted and filtered based on their abundance. Additionally, and contrary to other methods, during one of the index-building steps, for a given $k$-mer, its abundance in all datasets is known. This offers the possibility to conserve a $k$-mer having an abundance lower than the fixed threshold, which would be filtered out by other methods, but having some occurrences in the other datasets. This may reflect the presence in one of the samples of a low abundant species for which we want to preserve the data. Once $k$-mers are filtered, sub-matrices are built. Each sub-matrix indexes a subset of $k$-mers matching a specific set of minimizers.

As represented in Figure 1 (right), the resulting index built by `kmindex` consists of $P$ distinct matrices (with $P$ being the number of partitions, equal to 3 in the figure). To save indexing and query time, the index is "inverted": given a $k$-mer, the $N$ bits indicating its presence/absence in the $N$ indexed datasets are consecutive in the index. This allows for fast queries across numerous datasets. Hence, in practice, in a matrix, each row is a bit vector representing the presence or absence of a hash value in each indexed sample. Note that the rows are not packed to save construction and query time. This results in the fact that each row is composed of $\lceil \frac{N}{8} \rceil \times 8$ bits. Doing so, $\min(0, 8 - N \bmod 8)$ bits are unused for each row, as represented by a double arrow in Figure 1. This is up to 7 bits per row. These few lost bits may appear as a drawback, but this is negligible regarding the $N$ value that is meant to be in the order of a few hundred or thousands, and, importantly, this enables us to efficiently append novel indexed samples to an existing index.

By default, the resulting index is not compressed. Although requiring more space, this ensures optimal access time (both for writing and reading), and it offers the possibility to dynamically append new datasets to an existing index.

##### 1.2 Index query

The query process introduced in `kmindex` is also sketched in Figure 1. Batch processing is used for queries. This allows maximum throughput while maintaining control over memory usage. The user can specify the batch size and the maximum number of parallel batches according to the system's capabilities.

The resolution of a batch proceeds as follows.

1. **Bucketing**. The index is organized by partition, each corresponding to a set of minimizers. The first step consists of splitting query sequences into $k$-mers, which are then hashed and inserted into the

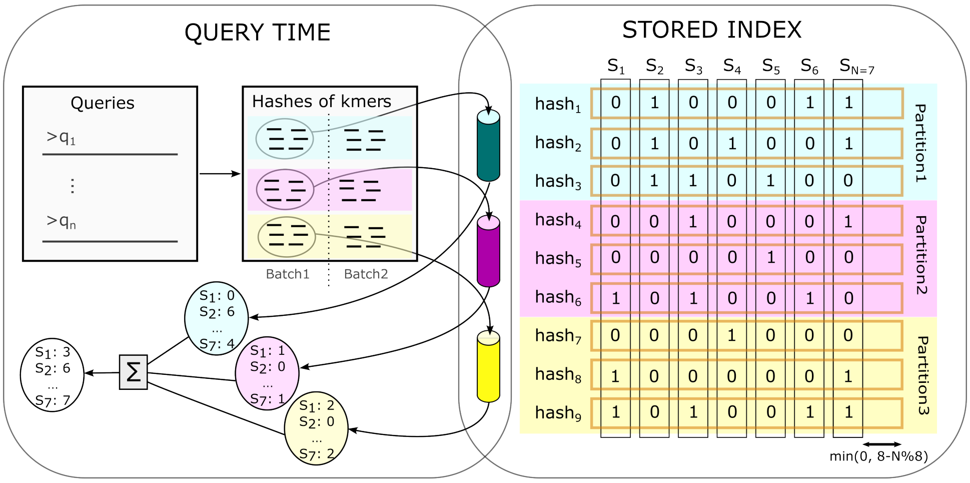


Figure 1: `kmindex`: an overview of the data structure and the query process. **Right: Data structure showing the stored index on disk**. Vertical black rectangles represent the $N$ Bloom filters (one per input sample). The orange horizontal rectangles represent the actual data structure saved on disk, storing consecutively the 0/1 values of distinct Bloom filters for the same hash value. The three colored horizontal rectangles represent three partitions, each saved in a distinct file. **Left: Query example**. Hashes of $k$-mers are symbolized by small horizontal lines, divided into two batches, and grouped by partitions. Each group is sorted. Each cylinder represents the streaming of a set of hashed $k$-mers, querying lines of Bloom filters mapped into memory. For simplicity, the image shows only queries from hashed $k$-mers from one of the two represented batches. In practice, for each partition, all batches are queried. For each sample, the results from each partition and each batch are finally summed up (as symbolized by the "Σ" symbol in this figure). Also for simplicity, this figure does not represent the use case in which the distribution of hits along the query sequences is reported as a binary vector.

right partition according to their minimizers. Each $k$-mer partition of the batch can then be solved by querying the corresponding index partition.

2. **Sorting**. Each partition is sorted to enable its resolution in a single sequential pass on the corresponding index partition, avoiding cache-misses.

3. ***k*-mer level resolution**. Querying a specific $k$-mer consists of fetching the row that corresponds to its hash value in the index to retrieve the bit vector corresponding to its presence or absence in each sample. For each query, the response vectors are aggregated by summation, resulting in an integer vector that represents the number of positive hits in each indexed sample.

   Obtaining the response vector for each $k$-mer is the current bottleneck because of I/O operations. For this reason, instead of loading the index into memory, index partitions are read through memory-mapped files. This allows reading only the parts of the index that are relevant to the batch resolution, which is particularly beneficial in the case of small queries.

   The memory-mapped files can be managed in two different ways: **Normal mode**: Each batch manages its own mappings of index partitions. The mapping of a partition is closed as soon as all $k$-mers belonging to the partition are resolved. The cached pages are then marked as available for eviction, resulting in lower memory usage (see results Section 2.5). **Fast mode**: All batches share the same mappings. This way, a larger number of pages are kept in the cache when conditions are favorable, i.e. without memory pressure, avoiding possible new I/O operations when solving the remaining batches. The memory usage may therefore seem high due to important page caching, up to the size of the index in the context of large query sets. Note that both modes require the same minimum amount of memory, the other part of the memory usage corresponds only to page caching which is automatically managed by the kernel. In other words, under memory pressure, both modes show the same memory usage.

4. **Sequence level resolution**. Finally, a result file is generated in either "json" or "tsv" format depending on the user's choice. Query results are filtered based on the threshold specified by the user. The user can also request the distribution of hits along the query sequences, represented as a binary vector.

##### 1.3 Drastically reducing the false positive rate using `findere`

The `kmindex` algorithm embeds the `findere` approach [10]. `findere` enables a drastic reduction of the false positive rate when querying successive $k$-mers from a query while using an Approximate Membership Query data structure (AMQ) (such as Bloom Filters) for indexing. For each indexed dataset, the central idea consists in indexing its $s$-mers instead of its $k$-mers in the AMQ, with $s \leq k$. At query time, a $k$-mer is considered as existing in the dataset if all its $k-s+1$ constituent $s$-mers are reported as present by the AMQ. In the general case, using this approach, a $k$-mer is wrongly reported as present (a false positive) when all its $s$-mers are themselves false positives. This has the effect of exponentially decreasing the false positive rate with respect to the $k-s$ value. When querying $k$-mers from a sequence of length $n$, in the general case, $n-k+1$ calls to the AMQ have to be made. Using `findere`, $n-s+1$ calls must be made ($k-s$ more than without using `findere`). This has negligible and no measurable impact on query time.

### 2 Benchmarks

#### 2.1 Setup

In situations involving large indexes and queries, various factors such as I/O operations or caching can impact performance. To account for these effects, we performed the benchmarks from a user's perspective. As a result, all measurements are obtained using the command line tools with particular attention to caching effects. The reported values include input parsing, query execution, and output writing. The results presented in Section 2.5 demonstrate the performance in a cold (the most likely and also the least favorable) or warm cache context.

We recall that the dataset for this benchmark is composed of metagenomic seawater sequencing data from 50 *Tara* Oceans samples, of 1.4TB of gzipped fastq files. It contains approximately 1,420 billion $k$-mers. Among them, approximately 394 billion are distinct, and 132 billion occur twice or more. These data are publicly available.

Executions were performed on the GenOuest platform on a node with 64 cores (128 threads) Xeon 2.2 GHz (L1 = 48KB, L2 = 1.25MB, L3 = 48MB shared) with 900 GB of memory. All computations are performed on an *xfs* filesystem allowing 1052MB/s sequential reads, 473MB/s sequential writes, and 908MB/s random reads (throughput measurements are obtained using `fio`: `https://github.com/axboe/fio`). All tested tools were parameterized to use 32 threads.

Used commands, data accession identifiers, input random sequence, and output files are provided in the `https://github.com/pierrepeterlongo/kmindex_benchmarks` companion website.

#### 2.2 Index construction

Indexes were constructed using $k = 28$. In the case of `kmindex`, which uses the `findere` approach, the 28-mers are emulated using $s$-mers of size $s = 23$ (see Section 1.3).

| | Wall clock time | Max Memory (GB) | Max temp. disk (GB) | Comment |
|---|---|---|---|---|
| `PAC` [8] | 14h59 | 190 | $191 + 1415^{\beta}$ | Empty result to queries |
| `ggcat` [3] | 12h59 | 69 | $2472 + 1415^{\beta}$ | ⚠Max No open files: 211,648. Query of one sequence killed after 12h. |
| `Themisto` [1] | 9h14 (killed) | > 900 | 4261 | Killed because of RAM usage |
| `HIBF` [9] | 16h53 (killed) | > 900 | 0.3 | Killed because of RAM usage |
| `MetaGraph` [6] | 50h53 (killed) | > 900 | 2144 | Killed because of RAM usage |
| `Bifrost` [5] | 12h57 (killed) | > 900 | 0 | Killed because of RAM usage |

$^{\beta}$ in order to consider multiple files per sample, the original input file has to be concatenated and so doubled using `PAC`, `ggcat`.

Table 1: Tested tools for which we were not able to build an index on the 50 *Tara* ocean samples or for which we were not able to perform a query. Tested tool versions: `PAC` commit cee1b5c (as used in the original PAC paper) and commit 940f18b (following a personal communication with the authors), `ggcat`: version v0.1.0, `Themisto`: version v3.2.0, `HIBF`: (`Raptor` version: 3.0.0, `Sharg` version: 1.0.1-rc.1, `SeqAn` version: 3.3.0-rc.1), `MetaGraph` version: 0.3.6., `Bifrost` version: 1.3.0.

Except for `kmindex`, only two tools, `MetaProFi` and `COBS`, were able to finish building the index and perform queries. Table 1 shows results for other tested tools that did not finish to build the index or for which the queries were not possible.

| | Step | Wall clock time | Max Memory (GB) | Max temp. disk (GB) | Output size on disk (GB) |
|---|---|---|---|---|---|
| MetaProFi [11] | KMC3 count | 3h44 | 278 | 1019 | 1019 |
| | KMC3 dump | 18h11 | 0 | 5684 | 5684 |
| | MetaProFi | 8h20 | 232 | 226 | 226 |
| | Overall | 30h15 | 278 | 5684 | 226 |
| COBS [2] | KMC3 count | 3h44 | 278 | 1019 | 1019 |
| | KMC3 dump | 18h11 | 0 | 5684 | 5684 |
| | COBS | 4h35 | 160 | 184 | 184 |
| | Overall | 26h30 | 278 | 5684 | 184 |
| kmindex | All | **2h56** | **107** | **878** | **164** |

Table 2: Comparing kmindex indexing performances to MetaProFi and COBS. The indexed dataset is composed of 50 *Tara* Oceans metagenomes datasets (total size 1.4TB). "Wall clock time" corresponds to the *user* time. "*Max temp. disk*" indicates the maximal disk used at runtime, this can be a temporary usage as for kmindex. "*Output size on disk*" indicates the size of the created files. kmindex can be run with a single command line, here resumed in the "*All*" step. Tested tool versions: kmindex version: 0.4.0, MetaProFi version 0.6.0, COBS commit 1cd6df2. Both COBS and MetaProFi require to filter $k$-mers upstream. This was performed using KMC3 [4], version 3.2.2.

Recall that count and filter $k$-mers are mandatory steps for all $k$-mer-based indexing tools. For example, 67% of the $k$-mers in this dataset are unique. Indexing them would more than double the index size and produce imprecise query replies. However, these time-consuming steps are not included in MetaProFi and COBS. This is why, for these two tools, and before constructing the indexes, we used KMC3 to count $k$-mers and remove those seen only once and thus considered as erroneous.

Table 2 shows that kmindex is ≈ 10 times faster and uses ≈ 3 times less memory and ≈ 6.5 less disk than the only other tools able to finish the index building with 900 GB of RAM and to perform a query in less than 12h. These results highlight the fact that time and memory usage are a bottleneck for tools that require to build a compacted version of the input data. Overall, for building the index, kmindex exhibits better performances on all considered criteria.

#### 2.3 Query performances

| No. queries | 1 | 10 | 100 | 1,000 | 10,000 | 100,000 | 1,000,000 | 10,000,000 |
|---|---|---|---|---|---|---|---|---|
| MetaProFi Time | 12s72 | 15s28 | 1m33 | 2m57 | 3m02 | 3m37 | 11m56 | 1h29 |
| MetaProFi Memory peak (GB) | 0.3 | 0.3 | 0.3 | 0.32 | 0.44 | 2.25 | 21 | 203 |
| COBS Time | 1s51 | 1s41 | 1s91 | 10s73 | 1m37 | 15m14 | 2h00 | 15h56 |
| COBS Memory peak (GB) | 0.012 | 0.018 | 0.036 | 0.28 | 2.66 | 24.58 | 138 | 295 |
| kmindex Time | **0s06** | **0s23** | **0s87** | **3s94** | **18s03** | **58s35** | **1m13s** | **4m21s** |
| kmindex Memory peak (GB) | **0.005** | **0.005** | **0.006** | **0.01** | **0.05** | **0.45** | **4.9** | **46.7** |

Table 3: Query time performance of the indexes on the 50 *Tara* Ocean samples. Queries are composed of reads uniformly sampled from the 50 *Tara* Oceans datasets. Executions were performed on a cold cache.

Table 3 details the results provided in the main text and proposes an extended range of results. Notably, it indicates the peak RAM usage at query time, not included in the main text. This highlights the limited RAM usage of kmindex compared to MetaProFi and COBS during queries.

#### 2.4 False positive rates

In order to test the FP rate, we generated a random sequence (25% chance of each nucleotide at each position, ≈50% GC) of size 10000. We used it for querying the index of the 50 *Tara* Oceans samples, successively querying the 9973 (10000-28+1) overlapping 28-mers of the query sequence. Note that we do not have a way to assess if each queried random $k$-mer occurs in the indexed set or not. Thus it may appear by chance that such a random $k$-mer indeed occurs in the set. This happens with a probability of $\times 10^{-9}$ in the biggest set. Hence the reported False Positive rate is an upper bound. This detail does not impact the conclusions offered by the results.

Results are presented in Table 4. As stated in the main manuscript, MetaProFi and COBS, the only tested tools with which we were able to perform queries, have an average false positive rate of 11.18% and 13.29%, respectively. In contrast, with a similar and smaller index size, kmindex shows a negligible average false positive rate of 0.006%.

| | Average | Median | Min | Max |
|---|---|---|---|---|
| Theoretical | 11.62 | 10.77 | 6.86 | 21.25 |
| `MetaProFi` | 11.18 | 9.92 | 6.93 | 21.55 |
| `COBS` | 13.29 | 12.30 | 7.07 | 24.60 |
| `kmindex` | **0.006** | **0** | **0** | **0.18** |

Table 4: False positive rates (in %). Indexed: 50 *Tara* Oceans samples. Queried: $k$-mers ($k = 28$) from a random sequence of size 10k nucleotides. Theoretical results correspond to the usage of BFs of size 30 billion bits as used by `MetaProFi`. The `COBS` index was built using the "-f 0.25" option to set the FP rate to 25%. `kmindex` results are below the expected theoretical rates at it implements the `findere` approach [10] (see Section 1.3).

#### 2.5 Warm and cold query results, and "fast-mode"

| | cache | | Querying 1 | 10 | 100 | 1k | 10k | 100k | 1M | 10M |
|---|---|---|---|---|---|---|---|---|---|---|
| **kmindex** | **c** | **T (s)** | 0.06 | 0.23 | 1.24 | 4.71 | 19.78 | 53.72 | 93.90 | 261 |
| | | **M (GB)** | 0.005 | 0.005 | 0.006 | 0.01 | 0.05 | 0.45 | 4.9 | 46.7 |
| | **w** | **T** | 0.06 | 0.20 | 1.15 | 4.02 | 10.84 | 16.42 | 40.76 | 225 |
| | | **M** | 0.005 | 0.006 | 0.006 | 0.01 | 0.06 | 0.43 | 4.70 | 42.6 |
| | **w+** | **T** | 0.02 | 0.1 | 0.74 | 2.65 | 5.64 | 13.54 | 42.12 | 227 |
| | | **M** | 0.005 | 0.006 | 0.006 | 0.01 | 0.05 | 0.44 | 4.48 | 43.70 |
| **kmindex fast** | **c** | **T** | 0.06 | 0.10 | 0.31 | 2.34 | 16.56 | 44.87 | 61.50 | 98.52 |
| | | **M** | 0.005 | 0.009 | 0.035 | 0.29 | 2.83 | 25.7 | 133 | 194 |
| | **w** | **T** | 0.03 | 0.08 | 0.24 | 1.57 | 7.20 | 7.86 | 15.79 | 64.36 |
| | | **M** | 0.005 | 0.009 | 0.035 | 0.29 | 2.84 | 25.7 | 133 | 194 |
| | **w+** | **T** | 0.06 | 0.05 | 0.07 | 0.18 | 1.04 | 4.63 | 15.18 | 62.33 |
| | | **M** | 0.005 | 0.009 | 0.035 | 0.29 | 2.84 | 25.7 | 133 | 194 |

Table 5: Time (seconds) and memory (GB) performances when querying from 1 read to 10 million reads over the 50 *Tara* Ocean samples indexed with `kmindex`. We consider the following scenarios: '**c**': cold (empty cache), '**w**' warm (successive distinct queries), and '**w+**' warm+ (successive identical queries). Queries are performed using 32 threads.

Table 5 shows time and memory usage results when performing queries using `kmindex`. We evaluate 3 query scenarios from the least to the most favorable: '**c**' cold (empty cache), '**w**' warm (successive distinct queries), and '**w+**' warm+ (successive identical queries). As expected, the query times decrease drastically with the cache benefit, illustrating the I/O bounds of `kmindex`. Note that the differences between '**w**' and '**w+**' decrease with the number of queries. Indeed, running a very large number of arbitrary queries increases the probability of loading useful pages for subsequent queries.

In *fast* mode, the kernel is allowed to keep as many pages as possible in the cache resulting in significantly faster queries at the cost of higher memory usage (near to the index size for 10 million queries). Under memory pressure, the memory usage would be equivalent to the *normal* mode.

#### 2.6 `kmindex` dynamicity performances

| No. queries | 1 | 10 | 100 | 1,000 | 10,000 | 100,000 | 1,000,000 | 10,000,000 |
|---|---|---|---|---|---|---|---|---|
| `kmindex` original (50 samples at once) | 0.13 | 0.13 | 0.40 | 2.46 | 16.52 | 41.44 | 54.87 | 100.36 |
| `kmindex` merge (5×10 samples merged) | 0.14 | 0.15 | 0.40 | 2.47 | 16.54 | 43.92 | 53.81 | 92.26 |
| `kmindex` register (5×10 samples registered) | 0.37 | 0.64 | 1.91 | 10.76 | 45.68 | 72.68 | 82.43 | 259.98 |

Table 6: Time performances (seconds) when querying from 1 read to 10 million reads over the 50 *Tara* Ocean samples indexed with `kmindex`. The index is built either as "**original**", "**merged**", or "**register**". With the "**original**" approach, the 50 samples are indexed in a unique process. With the "**merged**" and the "**register**" approaches, the 50 samples are separated into 5 groups of 10 samples each. The "**merged**" approach consists of physically extending an existing index, thus ending up with a unique index with the same performances as in the "**original**" approach. The "**register**" approach consists of registering independent indexes together.

`kmindex` disposes of two distinct ways to add novel samples to an existing index. The indexing time does not depend on the chosen approach. As presented in Table 6, when merging indexes together, the query time is optimal, equivalent to the one obtained from the same index built directly on the full dataset. However, this approach has the constraint that all the merged indexes have to be built using the same parameters (hash function, number of partitions, bloom filter sizes). On the other hand, when distinct indexes are *registered* together, each index is individually queried increasing the running time. However, as the indexing parameters are independent, this solution is more flexible. It is well adapted when indexing highly diverse samples such as samples from other *Tara* missions or distinct phylogenetic groups for instance.

### 3 The Ocean Read Atlas

#### 3.1 Dataset

ORA index is composed of 1,393 samples (distinct locations and distinct fraction sizes) of the *Tara* Oceans project. These samples are divided into six distinct groups, determined by the size fraction of the sequenced species. These fractions correspond to the physical filter sizes used during the sampling campaign. Based on this clustering we built six distinct indexes (all with the same parameters). At query time, as all the six indexes are registered in a unique meta-index, the whole set of samples is queried. A description of the dataset is available in Table 7. The size of the final uncompressed index is approximately 13% of the size of the raw fastq.gz files, which is 36.7 TB.

| Fraction size | Number of samples | Average number of distinct $k$-mers per sample |
|---|---|---|
| Integrated ($0.8-2000\mu m$) | 193 | 14.5e9 |
| Meso ($180-2000\mu m$) | 208 | 13.2e9 |
| Micro ($20-280\mu m$) | 195 | 15e9 |
| Nano ($3-20\mu m$) | 213 | 16.1e9 |
| Pico ($0.2-5\mu m$) | 425 | 11.1e9 |
| Virus ($< 0.2\mu m$) | 159 | 2.3e8 |

Table 7: Description of the indexed dataset organized by size fraction. The "Fraction size" column indicates the size range of the target sequenced species.

##### 3.1.1 Sequencing data availability

Shotgun metagenomic sequences of all the samples from the *Tara* Oceans Expedition (2009–2013) are available at the European Nucleotide Archive (ENA, `https://www.ebi.ac.uk/ena/`) under global accession number PRJEB402 (PRJEB1787 and PRJEB9740 for bacteria and archaea, PRJEB1788 for giant viruses, PRJEB4352 and PRJEB9691 for protists, and PRJEB4419 and PRJEB9742 for DNA viruses.

##### 3.1.2 Environmental data

The environmental data are from the *Tara* Oceans Expedition (2009–2013) and are available on PANGAEA (`https://doi.pangaea.de/10.1594/PANGAEA.875582`). The environmental database contains currently:

- BIODIV: `https://doi.org/10.1594/PANGAEA.853809`
- CARB: `https://doi.org/10.1594/PANGAEA.875567`
- HPLC: `https://doi.org/10.1594/PANGAEA.875569`
- MESOSCALE: `https://doi.org/10.1594/PANGAEA.875577`
- NUT: `https://doi.org/10.1594/PANGAEA.875575`
- SENSORS: `https://doi.org/10.1594/PANGAEA.875576`
- SEQUENCING: `https://doi.org/10.1594/PANGAEA.875581`
- WATERCOLUMN: `https://doi.org/10.1594/PANGAEA.875579`

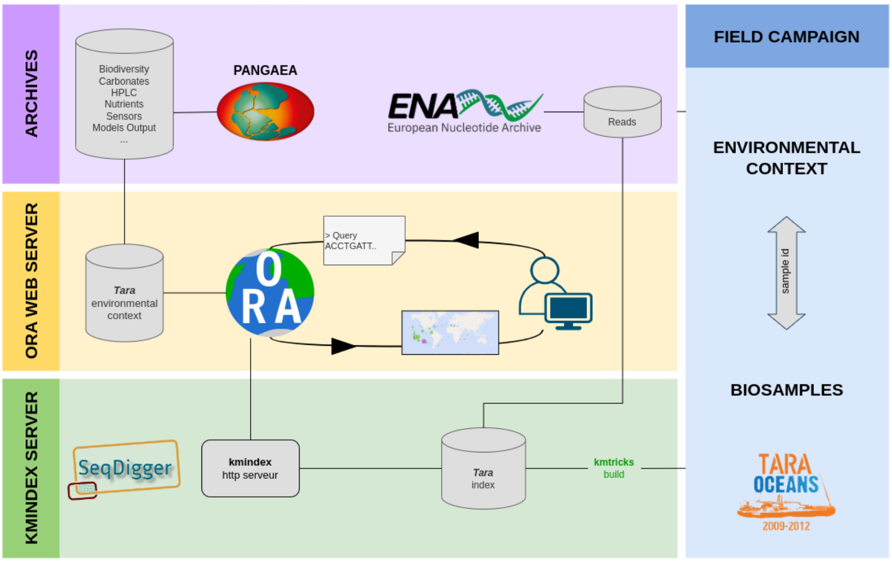


Figure 2: The ORA web service is organized in 3 distinct entities: **1)** a database containing the environmental parameters and the biosample information of the campaign, **2)** the `kmindex` server allowing index request, and **3)** the ORA server making the link between the 2 previous entities and allowing visualization of the results *via* a web interface.

#### 3.2 Workflow

The Ocean Read Atlas (ORA) service is composed of 3 parts:

1. the environmental database containing the parameters measured and estimated during the sampling of the *Tara* Oceans Expedition;

2. the `kmindex` server to query the `kmindex` index made with sequencing reads from *Tara* Oceans Expedition *via* HTTP requests;

3. and the ORA server to query the index and make the link between $k$-mers (contained in the query sequence) and the environmental parameters of samples sharing these $k$-mers. ORA provides results on a webpage including maps, plots, and table files.

A representation of these components is available in Figure 2.

#### 3.3 `kmindex` server

The index is stored on a Ceph storage cluster (SSD pool). The CRUSH algorithm enables the Ceph storage cluster to scale, rebalance, and recover data dynamically. Nodes of this storage cluster are on a redundant network using LACP trunking configuration.

The `kmindex` HTTP server runs in a Qemu/KVM virtual machine (VM) with 16 cores and 32 GB RAM, supporting 16 concurrent queries. VM is stored on a Proxmox virtualization cluster with HA capabilities. Depending on our available resources, storage, and VM capacities may be expanded if needed. This infrastructure is mandatory to ensure service continuity.

Figure 2 represents the overall `ORA` workflow.

#### 3.4 About `ORA` usages and limitations

The service currently supports unique query of a FASTA file limited to 10 kilobase pairs *via* the web interface. Each user is limited to 200 jobs per 24 hours. The results can either be delivered directly or sent by email with a link valid for 2 weeks. In the future and depending on our computational capacity, we expect to offer an API with more features such as the integration of the query abundance and metadata associated with target sequences.

A user guide manual is available at the following address: `https://ora.mio.osupytheas.fr/manual/pages/`. In the interfaces section (`https://ora.mio.osupytheas.fr/manual/pages/interfaces.html`)

### 4 Code and reproducibility

- `kmindex` source code: `https://github.com/tlemane/kmindex/`
- `kmindex` documentation: `https://tlemane.github.io/kmindex/`
- `ORA`: `https://ora.mio.osupytheas.fr/`
- `ORA` source code: `https://gitlab.osupytheas.fr/ocean_atlas/ora/`
- Benchmark data description and used commands: `https://github.com/pierrepeterlongo/kmindex_benchmarks/`

[6] Mikhail Karasikov, Harun Mustafa, Daniel Danciu, Marc Zimmermann, Christopher Barber, Gunnar Rätsch, and André Kahles. Metagraph: Indexing and analysing nucleotide archives at petabase-scale. *BioRxiv*, 2020.

[7] Téo Lemane, Paul Medvedev, Rayan Chikhi, and Pierre Peterlongo. kmtricks: Efficient and flexible construction of bloom filters for large sequencing data collections. *Bioinformatics Advances*, 2022.

[8] Camille Marchet and Antoine Limasset. Scalable sequence database search using Partitioned Aggregated Bloom Comb-Trees. In *Recomb 2022- 26th Annual International Conference on Research in Computational Molecular Biology*, La jolla, United States, May 2022.

[9] Svenja Mehringer, Enrico Seiler, Felix Droop, Mitra Darvish, René Rahn, Martin Vingron, and Knut Reinert. Hierarchical interleaved bloom filter: enabling ultrafast, approximate sequence queries. *Genome Biology*, 24(1):1–25, 2023.

[10] Lucas Robidou and Pierre Peterlongo. findere: fast and precise approximate membership query. In *International Symposium on String Processing and Information Retrieval*, pages 151–163. Springer, 2021.

[11] Sanjay K Srikakulam, Sebastian Keller, Fawaz Dabbaghie, Robert Bals, and Olga V Kalinina. Metaprofi: an ultrafast chunked bloom filter for storing and querying protein and nucleotide sequence data for accurate identification of functionally relevant genetic variants. *Bioinformatics*, 39(3):btad101, 2023.
